# NAD precursors cycle between host tissues and the gut microbiome

**DOI:** 10.1101/2021.11.15.468729

**Authors:** Karthikeyani Chellappa, Melanie R. McReynolds, Wenyun Lu, Xianfeng Zeng, Mikhail Makarov, Faisal Hayat, Sarmistha Mukherjee, Yashaswini R. Bhat, Siddharth R. Lingala, Rafaella T. Shima, Hélène C. Descamps, Timothy Cox, Lixin Ji, Connor Jankowski, Qingwei Chu, Shawn M. Davidson, Christoph A. Thaiss, Marie E. Migaud, Joshua D. Rabinowtiz, Joseph A. Baur

## Abstract

Abstract

Nicotinamide adenine dinucleotide (NAD) is an essential redox cofactor in both mammals and microbes. Here we use isotope tracing to investigate the precursors supporting NAD synthesis in the gut microbiome. We find that preferred dietary NAD precursors are absorbed in the proximal part of the gastrointestinal tract and not available to microbes in the distal gut. Instead, circulating host nicotinamide enters the gut lumen and supports gut microbiome NAD synthesis. In addition, the microbiome converts nicotinamide, originating from the host circulation, into nicotinic acid. Host tissues uptake and utilize this microbiome-derived nicotinic acid for NAD synthesis, maintaining circulating nicotinic acid levels even in the absence of dietary consumption. Moreover, the main route from oral nicotinamide riboside, a widely used nutraceutical, to host NAD is via conversion into nicotinic acid by the gut microbiome. Thus, NAD precursors cycle between the host and gut microbiome to maintain NAD homeostasis.

## Introduction

Nicotinamide adenine dinucleotide (NAD) is an indispensable reduction-oxidation (redox) co-enzyme. It is also a substrate for signaling enzymes including sirtuins, poly-ADP polymerases (PARPs), CD38 and SARM1. NAD-consuming enzymes degrade the NAD molecule and release nicotinamide (NAM), thus necessitating a balance between synthesis and consumption to maintain NAD homeostasis. NAD levels have been reported to decline in a spectrum of pathological conditions including aging, obesity, inflammatory, cardiovascular, and neurodegenerative diseases. In many cases, the decline in cellular NAD content correlates with mitochondrial and metabolic dysfunction. NAD boosting ameliorates metabolic perturbations in rodent models of disease^1–5^ and has shown promise in some human clinical trials^6, 7^, albeit with modest or no effects in others^8–10^.

Animals have coevolved with commensal microorganisms, including the gut microbiome. The microbes that dwell in the gut lumen are unique in that they can potentially access both ingested food and circulating nutrients to support their metabolic activity. That said, nutrients that are absorbed in the proximal parts of the small intestine are largely inaccessible to the microorganisms residing in the gut lumen. Moreover, it is unclear which, if any, circulating host nutrients permeate the gut microbiome sufficiently to drive microbiome metabolism. Two established nutrient sources of the colonic microbiome are poorly digestible complex carbohydrates (fiber)^11^ and host mucus^12–17^.

As with host cells, NAD is essential to microorganisms. Tremendous effort has been undertaken over the past century to understand NAD metabolism in mammalian tissues and free-living microbes. In contrast, NAD metabolism in the gut microbiome is less extensively studied. A recent study revealed a profound decrease in the NADH/NAD redox ratio (but not absolute NAD levels) in colonocytes of germ-free mice^18^. Compared to conventional mice, germ free mice fed a western diet had higher liver NAD^19^. Intriguingly, gut microbiota are required for the full NAD-boosting effect of orally delivered precursors including both NAM and the common nutraceutical nicotinamide riboside (NR)^20^.

In mammals, three major routes of NAD biosynthesis are (1) *de novo* synthesis from tryptophan, (2) salvage synthesis from NAM via nicotinamide phosphoribosyltransferase (NAMPT), and (3) Preiss-Handler synthesis from nicotinic acid (which is also called “niacin”) by nicotinic acid phosphoribosyltransferase (NAPRT). *De novo* synthesis of NAD from tryptophan is quantitatively important mainly in the liver^21^, but the kidney and some immune cells also utilize this pathway to make NAD^22, 23^. Turnover of NAD generates NAM, which is exchanged with the systemic circulation. This provides a common pool of precursor that can be used to generate NAD through the salvage pathway. Nicotinic acid is obtained from the diet or microbiome, circulates at lower concentrations, and makes a quantitatively smaller contribution to NAD synthesis in most tissues^21^. NAD synthesis can be increased by exogenous supplementation with intermediates such as NR or NMN^1, 5, 24–26^. The degree to which this reflects direct incorporation of the parent molecule vs. breakdown and resynthesis is an active area of investigation^21^.

Most microbial species encode one or more enzymes required to synthesize NAD either *de novo* from amino acids or through salvage of “vitamin B3” (a collective term for nicotinic acid, NAM and related derivates such as nicotinamide riboside and nicotinic acid riboside)^27–30^ (Figure 1A). In unicellular microorganisms, *de novo* synthesis can occur either from tryptophan (as in mammals) or more commonly from aspartate by L-aspartate oxidase (*nadB*)^31^. Importantly, many bacteria (but not mammals) encode the enzyme to convert NAM to nicotinic acid: nicotinamide deamidase (*pncA*)^20, 27, 32^.

**Figure 1.**
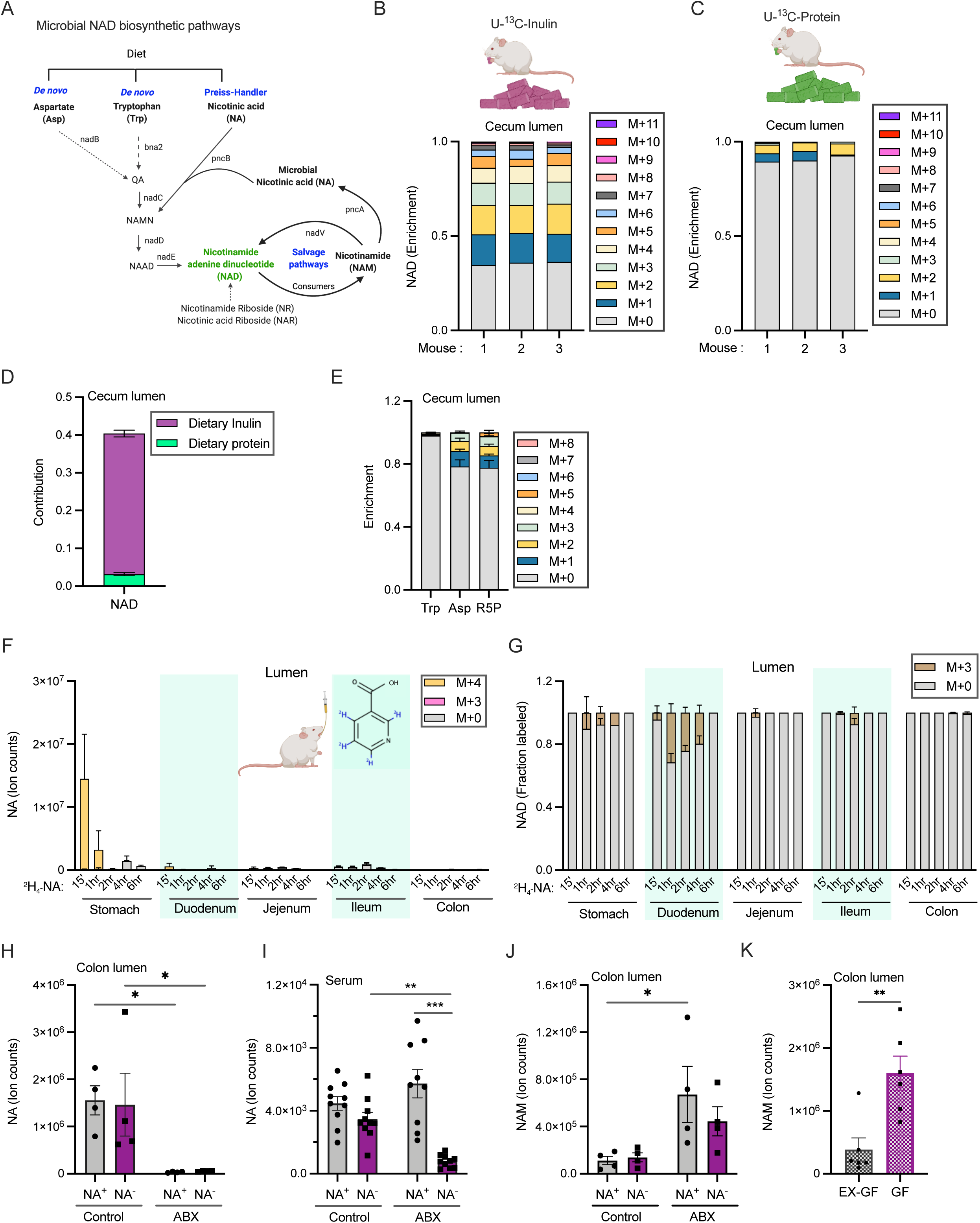
The majority of microbial NAD synthesis is not accounted for by dietary precursors. A. Different pathways used for NAD biosynthesis in microbes. B. Enrichment of carbon labeled NAD in the cecal lumen of mice fed with U-^13^C-inulin for 24h. n=3 mice per group. C. Enrichment of carbon labeled NAD in the cecal lumen of mice fed with U-^13^C-protein for 24h. n=3 mice per group. D. Contribution of U-^13^C-protein and U-^13^C-inulin to NAD synthesis in the cecal lumen of mice treated as in B and C. E. Enrichment of carbon labeled tryptophan, aspartate, and ribose phosphate in cecal lumen of mice treated as in B. F. Labeled and unlabeled NA content and fraction NAD (B) in the gut lumen after oral gavage of 1.96 μmoles of [2,4,5,6-^2^H]-NA, equivalent to one-third daily dietary nicotinic acid intake. G. Fraction labeled NAD in the gut lumen of mice treated as in F. H. NA content in the colonic lumen of conventional mice fed diet with (NA^+^) or without nicotinic acid (NA^-^) that were on drinking water or antibiotics cocktail (ABX). n=4 per group. I. NA content in the serum of mice treated as in H. n=9-10 mice per group. J. NAM content in the colonic lumen of mice treated as in H. n=4 per group. K. Abundance of NAM in the colonic lumen of germ-free (GF) and germ-free mice colonized with microbiota from specific pathogen free mice (Ex-GF). n=4 per group.

In this study, we use isotopic tracers to determine how the microbial NAD pool is established. We find that dietary protein and vitamin B3 are not primary precursors for NAD biosynthesis in gut microbiota. Instead, NAM in the host circulation enters the gut lumen to feed NAD biosynthesis, with fiber catabolism also feeding into NAD in the colonic microbiome. Beyond consuming host-derived NAM, the microbiome converts NAM into nicotinic acid, which provides an alternative NAD precursor to host tissues, restoring NAD levels in intestinal cells when salvage synthesis of NAD from NAM is blocked. The gut microbiome also converts orally delivered NR into nicotinic acid, which circulates at high levels post oral NR and is critical to NR’s capacity to boost host NAD. In short, we identify a NAM-nicotinic acid metabolic cycle that allows sharing of NAD precursors between host and microbiome.

## Results

### Soluble fiber contributes to microbiome NAD synthesis in the large intestine

We explored whether dietary fiber or protein provide precursors for *de novo* NAD biosynthesis by the gut microbiome. Grain-based chow contains around 15-20% of insoluble fiber (generally not metabolized) and 3-5% soluble fiber^33^ (fermented by colon bacteria)^27, 34, 35^. We tested whether ^13^C-labeled soluble fiber (inulin) or protein feed microbiome NAD synthesis. Mice were fed a refined diet mixed with 5% unlabeled inulin w/w for 14 days and then switched to diet with U-^13^C-inulin for 24h before sacrifice. Alternatively, they were fed standard chow containing U-^13^C-labeled protein for 24h. We found ∼65% of NAD molecules contained one or more labeled carbon atoms derived from inulin, versus less than 10% from protein (Figures 1B and 1C). Quantitative analysis accounting for precursor (e.g., amino acid) labeling in the cecum lumen and all carbon atoms in NAD revealed that fiber accounts for 40% of the total carbon contribution to NAD synthesis, versus only 3% for protein (Figures 1D and 1E, Supplementary Figure 1A). The contribution of inulin to microbial NAD synthesis was lower in other parts of the gut lumen and intestinal tissues (Supplementary Figure 1B). Fiber’s contribution to microbiome NAD included labeling of the key NAM moiety, as shown by MS/MS. Consistently, labeling was also detected in luminal NAM (Supplementary Figures 1C and 1D).

Simple sugars such as fructose are normally absorbed in the proximal part of the GI tract but may reach distal parts of the gut lumen under conditions of excessive nutrient intake^36^. We found that a large bolus of fructose supplied precursors for NAD synthesis by large intestinal microbiota (Supplementary Figures 1E-1J). Thus, carbohydrate that reaches the colon will feed into *de novo* NAD synthesis, but a large portion of NAD synthesis remains unaccounted for after considering dietary protein and fiber.

### Dietary nicotinic acid is not available to microorganisms residing in the distal gut

Standard mouse chow contains nicotinic acid, which could provide a precursor for NAD synthesis by gut microbes. To examine this possibility, we gavaged mice with labeled nicotinic acid (2,4,5,6-^2^H-NA) at a dose equivalent to one-third of daily intake. High levels of fully labeled nicotinic acid (M+4 NA) were detected in the stomach lumen at the earliest time point assayed (Figure 1F). Labeled NA in the serum peaked at 15 min and gradually declined (Supplementary Figure 1K). Incorporation of this tracer into NAD results in loss of label from the redox-active (4) position to generate M+3 metabolites. A small amount of unmetabolized M+4 NA was found in the lumen of the duodenum and jejunum, whereas only M+3 NA was detected in the distal small intestine and colon. While oral NA made a significant contribution to microbial NAD synthesis in the duodenal lumen (∼32%), the contribution was negligible in distal small intestine and large intestine (Figure 1G). These data are consistent with oral NA being absorbed or metabolized before reaching the lumen of the ileum and colon.

To establish the relative contribution of microbes to NA in the gut lumen and circulation, we next fed mice a refined diet with or without NA. Mice on each of the diets were either given normal drinking water or water treated with a cocktail of antibiotics to deplete microbiota. While antibiotics alone were sufficient to deplete NA in the gut lumen, a combination of NA-deficient diet and antibiotics was required to decrease circulating NA (Figures 1H and 1I).

### Circulating nicotinamide is a major precursor for luminal NAD and nicotinic acid

Strikingly, while antibiotics depleted NA in the gut lumen, they augmented luminal NAM (Figure 1J). A similar decrease in luminal NA and increase in NAM were observed in the intestinal lumens of germ-free mice (Figure 1K, Supplementary Figures 1L and 1M). These data raise the possibility that there is a microbiome-independent source of gut lumen NAM, and that microbes convert this NAM into NA.

What is the source of gut lumen NAM? One possibility is the flux of circulating NAM made by the host^37, 38^. To explore this possibility, we intravenously injected mice with 2,4,5,6-^2^H-NAM (Figure 2A). We detected fully labeled NAM in the lumen all along the GI tract (Figure 2B). In addition, M+4 labeled NA was found in the luminal contents along the length of the GI tract (Figure 2C). NA generated from infused NAM reenters the circulation as evidenced by labeled NA in serum (Figure 2D). Labeled NAD molecules were detected in the luminal contents of both the small and large intestines, indicating incorporation into the microbial pools (Supplementary Figure 2A). Similarly, intravenously delivered NAM tracer accumulated in the GI lumens of germ-free mice (Figure 2E and Supplementary Figure 2B) but was not converted into NA and NAD (Figures 2E and 2F, Supplementary Figures 2C and 2D).

**Figure 2.**
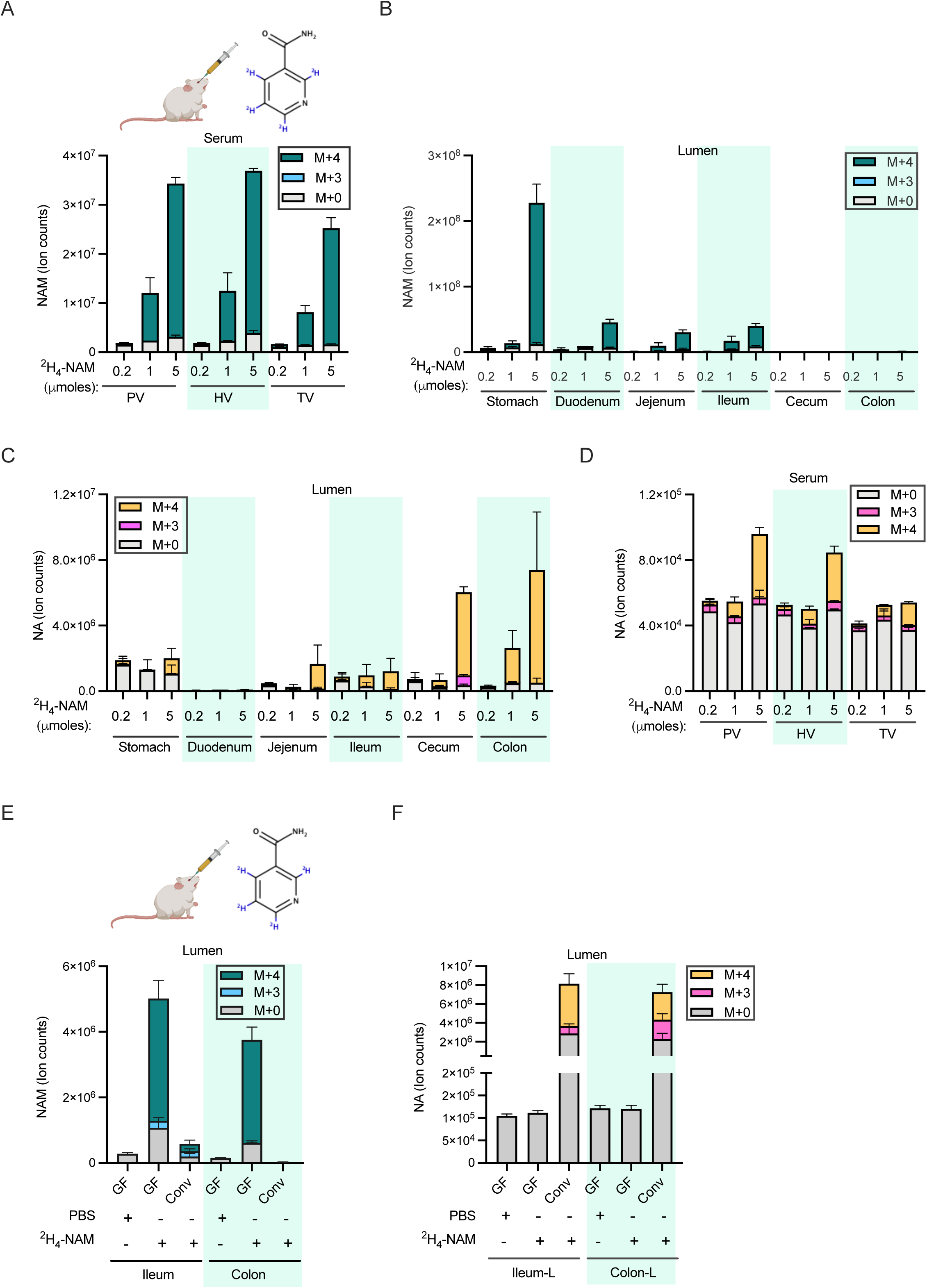
Circulating NAM enters the gut lumen and supports microbial NAD biosynthesis. A. NAM levels in the serum samples collected from portal (PV), hepatic (HV) and tail vein (TV) 15 min after retro-orbital injection of different doses of [2,4,5,6-^2^H]-NAM. n=2-3 mice per group. B. Labeled NAM detected in the lumen samples collected from mice treated as in A. C. Labeled and unlabeled NA detected in the luminal samples collected from mice treated as in A. D. Labeled and unlabeled NA detected in the serum samples collected from mice treated as in A. E. Labeled and unlabeled NAM in luminal samples collected from germ-free (GF) and conventional (Conv) mice retro-orbitally injected with 5 μmoles of [2,4,5,6-^2^H]-NAM and sacrificed after 2h. n=4 mice per group. F. Labeled and unlabeled NA in the luminal samples collected from mice treated as in E.

To rule out that the above results are due to large transient perturbations in circulating NAM levels following intravenous injection and to enable more quantitative downstream analyses, we next carried out experiments involving continuous intravenous infusion of 2,4,5,6-^2^H-NAM at a minimally perturbative rate (0.2 nmol/g/min). NAM infusion resulted in a rapid increase in serum M+4 NAM labeling and slower accumulation of circulating M+3 NAM derived from cycle(s) of NAD synthesis and breakdown in tissues (Figure 3A)^21, 39^. Both infused (M+4) and recycled (M+3) NAM were detected in the gut lumen, along with luminal and serum M+4 and M+3 NA (Supplementary Figures 3A-3C). NAD was labeled throughout the small intestine and colon, reaching steady-state in the ileum and jejunum lumen within 12h (Figure 3B) and colon at 18h. Despite higher uptake of NAM, labeling of NAD was slow in the stomach lumen. By normalizing to fractional NAM labeling in serum, we estimated the quantitative contribution of circulating NAM to microbiome NAD synthesis in different parts of the gastrointestinal tract (Figure 3C). Circulating NAM accounts for 80-90% of microbial NAD synthesis in the ileum and jejunum, and 45% in the colon (consistent with the colon microbiome synthesizing about half of its NAD *de novo* from fiber). Thus, circulating host NAM enters the gut lumen and is used by the microbiome for both NAD and NA synthesis.

**Figure 3.**
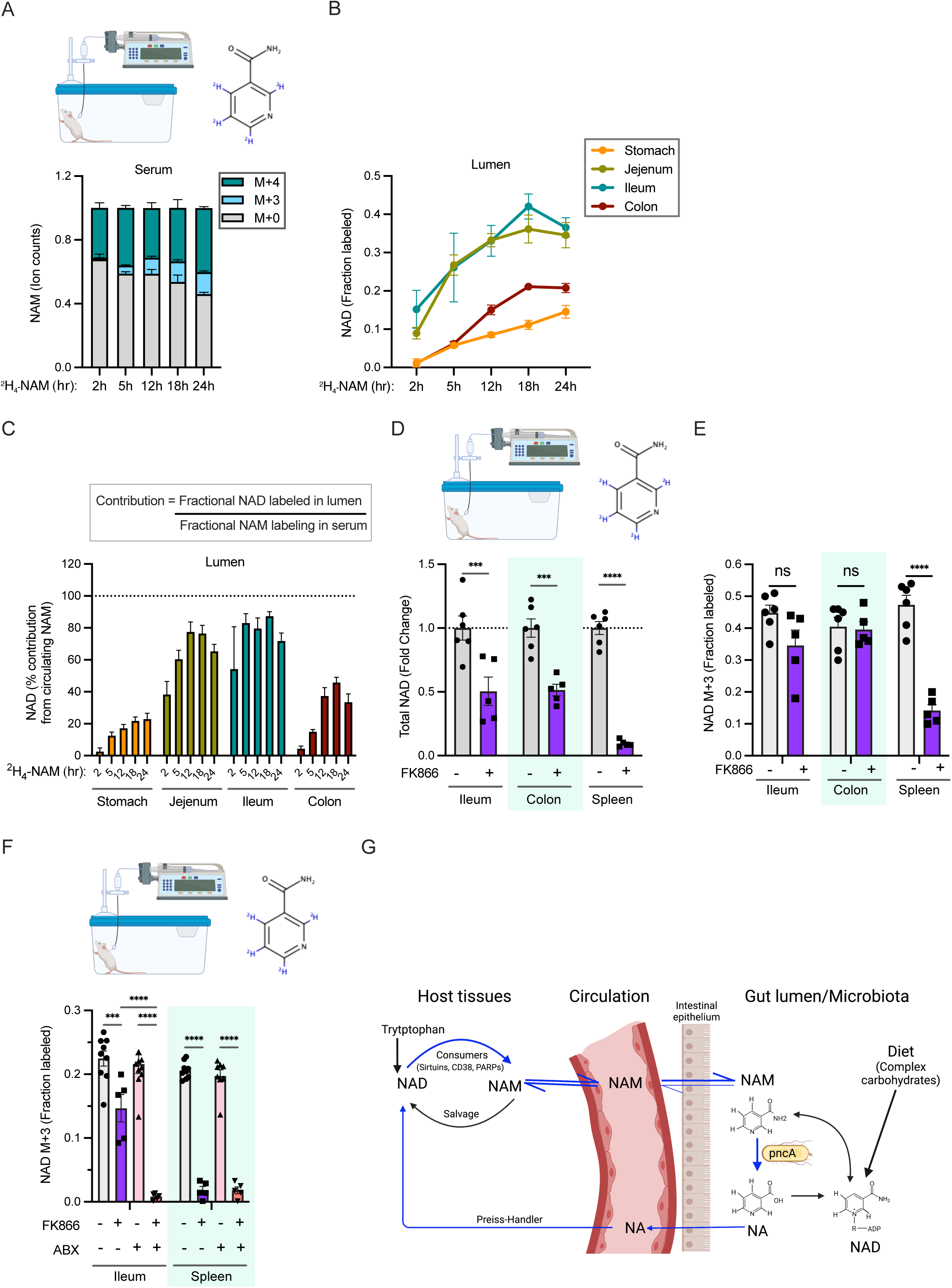
Vitamin B3 cycles between host and microbiome. A. Fraction labeling of nicotinamide in the serum of mice infused with 4mM [2,4,5,6-^2^H]-NAM for different time points. n=2-4 mice per group. B. Fraction labeled NAD in the luminal content of mice infused as in A. C. Percent contribution of circulating NAM to NAD synthesis in different parts of the gastrointestinal tract. D. Relative levels of total NAD in mice intraperitoneally injected with vehicle or FK866 and infused with [2,4,5,6-^2^H]-NAM for 24h. n=5-6 mice per treatment group. E. Fraction labeled NAD in mice treated as in D. Data for spleen in D and E was previously reported in McReynolds et al. Cell system, 2021. F. Fraction labeled NAD in tissues collected from control and antibiotics (ABX) treated mice intraperitoneally injected with vehicle or FK866 and infused with [2,4,5,6-^2^H]-NAM for 5h. n=3-5 mice per treatment group. G. Model showing a vitamin B3 cycle between host and microbes. Host-derived NAM in the circulation enters the gut lumen and is deamidated to NA by the microbiome. In turn, host tissues use microbiome-produced NA for synthesis of NAD, which is turned over to release NAM.

### Microbial deamidation bypasses salvage synthesis in host intestine and liver

Most tissues rely on the salvage pathway for NAD synthesis^21^. Inhibiting NAMPT, the rate-limiting enzyme in the salvage pathway, general depletes NAD levels in tissues in proportion to their NAD consumption rates^39^. Intestine is an exception to this trend. For example, intestine and spleen have similarly rapid NAD turnover rates, but treatment with the NAMPT inhibitor FK866 depletes >90% of NAD in spleen, but only about ∼50% in intestine^39^ (Figure 3D). Using 2,4,5,6-^2^H-NAM tracer, we observed persistent labeling of intestinal (but not spleen) NAD from the infused NAM even in the presence of NAMPT inhibitor (Figure 3E). We hypothesized that this persistent synthesis reflects intestinal incorporation of NA made by microbial deamidation of NAM, and thus would be blocked by a combination of NAMPT inhibitor and antibiotics. Indeed, FK866 treatment dramatically decreased NAD labeling from NAM in antibiotics treated mice (Figure 3F). In the liver, treatment of mice with either FK866 or antibiotics decreased labeling of NAD from infused labeled NAM, and combined treatment completely prevented intravenous NAM from contributing to hepatic NAD (Supplementary Figure 3D). Thus, gut microbiota provide an alternative route from NAM to NAD, bypassing the salvage synthesis pathway, in intestine and liver (Figure 3G).

### Nicotinamide riboside boosts tissue NAD pools via breakdown and deamidation to nicotinic acid

Nicotinamide riboside (NR) is widely promoted and consumed as an NAD-boosting nutraceutical supplement^40^. Intact NR can be taken up by cells and converted to NAD without the need for the energetically expensive endogenous metabolite phosphoribosyl pyrophosphate or the key salvage enzyme NAMPT. However, the majority of orally delivered NR has been shown to be rapidly cleaved to NAM before entering circulation and peripheral tissues^21, 41^. Moreover, oral NR supplementation has recently been shown to be much less effective at boosting liver and intestinal tissue NAD levels in the absence of gut microbiota, consistent with its being incorporated via a deamidated route^20^. We measured circulating levels of vitamin B3 species in mice following NR oral gavage (500 mg/kg). Circulating NA increased >100-fold increase, peaking at 50 μM. NAM showed a similar trend, with more extended exposure duration and modestly higher peak concentration (Figures 4A and 4B). The observed levels of NA from NR gavage are comparable to those in humans receiving therapeutic NA supplementation known to elicit effects on plasma lipids and lipoproteins^42–44^ and in excess of the EC_50_ for activation of GPR109A (the “niacin receptor”)^45–47^.

**Figure 4.**
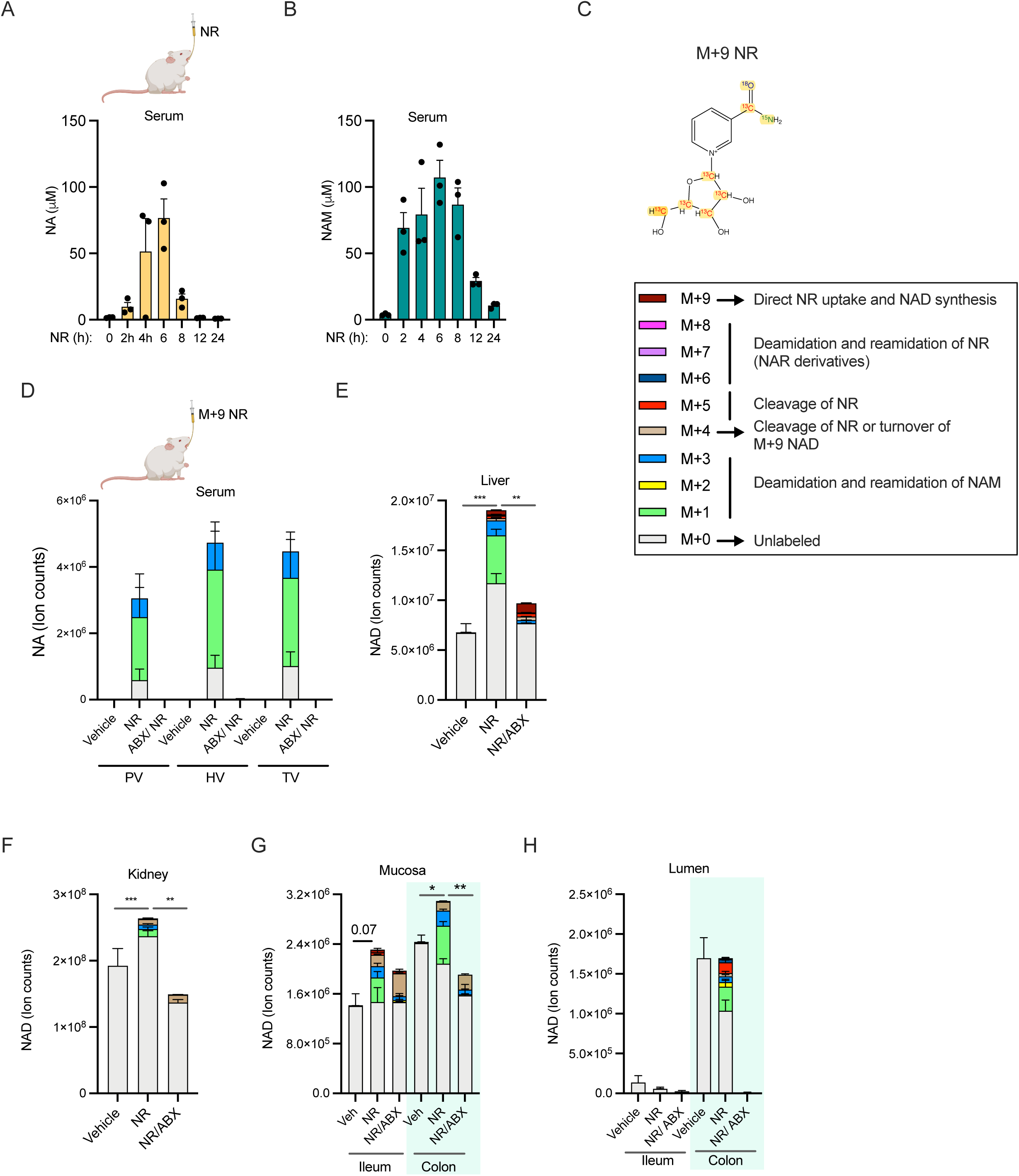
Oral NR supports NAD synthesis in mammalian tissues mainly via conversion to NA. A. Serum concentration of NA in mice orally gavaged with NR at a dose of 500mg/kg body weight n=3 per group. B. Serum NAM concentration in mice treated as in A. C. M+9 NR isotopically labeled in the amide group of nicotinamide moiety and ribose ring to establish the molecular fate of NR in host and microbes. D. Relative level of NA in the serum of control and antibiotics (ABX) treated mice orally gavaged with PBS (vehicle) or NR at a dose of 600 mg/kg body weight. Serum samples were collected from portal vein (PV), hepatic vein (HV) and tail vein (TV) after 3h of treatment (n=4-5 per treatment group). E. NAD isotopologues detected in the liver of mice treated as in D. F. NAD isomers in the kidney of mice treated as in D. G. Isotopologues of NAD in the intestinal tissues of mice treated as in D. H. Labeling pattern of NAD in the luminal samples collected from mice treated as in D.

Production of NAD and NA from NR can occur multiple ways, e.g., (i) NR to NAD to NAM to NA; (ii) NR to NAM to NA to NAD; (iii) NR to nicotinic acid riboside (NAR) to NAD and/or NA. Given that NR has been reported to be more effective than equivalent doses of NAM or NA^48^ and that metabolism of oral nicotinamide mononucleotide was recently suggested to proceed via the deamidated mononucleotide^49^, we were particularly intrigued by possibility (iii). To distinguish these routes, we synthesized an M + 9 isotopologue, on which ^15^N, ^13^C and ^18^O atoms have been introduced on the amide group in the NAM moiety and the five carbon atoms of the ribose ring are ^13^C (Figure 4C). Deamidation always removes the labeled nitrogen from the amide group and reamidation has a 50% chance of removing the labeled oxygen as well^50^. Orally delivered NR was found in the GI tract but not in other peripheral tissues, indicating rapid metabolism of NR before it enters the circulation (Supplementary Figure 4A). Only a small fraction of NAD in the liver and small intestine were M+9, indicating minimal direct assimilation of NR (Supplementary Figure 4B). Thus, route (i) is a minor contributor.

In both routes (ii) and (iii), the nicotinamide moiety of NAD is expected to be labeled as a mixture of M+1 and M+3, indicating synthesis via a deamidated species: NA or NAR. The routes can be distinguished because (iii) retains the original labeled ribose moiety, which would result in production of M+6 and M+8 NAD. Only M+1 and M+3 labeled NAD molecules were detected in the gut microbiome and host tissues (Supplementary Figure 4B), indicating separation of the NAM/NA moiety from the labeled ribose: assimilation via route (ii). Consistent with this, we detected M+1 and M+3 labeled nicotinic acid adenine dinucleotide (NAAD) in tissues (Supplementary Figure 4C). As expected for NR assimilation via a deaminated species, depletion of gut microbiota with antibiotics completely eliminated NR-derived NA from serum and dramatically abrogated the effect of NR supplementation on NAD in host tissues (Figures 4D-4H, Supplementary Figures 4D-4G). Direct incorporation of NR into NAD was higher in antibiotics-treated mice, but NAD synthesis from this route was not sufficient to significantly increase the total NAD pool in the absence of gut microbiota. Thus, NR is catabolized to NA by gut microbiota, and the resulting NA supports NAD synthesis in host tissues.

## Discussion

Hosts and their gut microbiomes have coevolved to have complex symbiotic relationships. While it is widely perceived that microbes extend digestive capabilities and produce metabolites such as short-chain fatty acids, flux of metabolites from host to microbiome remains limited to a few examples. These include the feeding of microbes on host-derived mucus^12–16, 51^, incorporation of urea nitrogen into microbial protein^52–54^, and bidirectional transport of lactate^55^. In addition, host-derived bile acids are metabolized by microbes and have a suppressive effect to limit microbiome growth^56–58^. In each of these cases, metabolism of host molecules by the microbiome may be considered opportunistic, even if it is ultimately mutually beneficial. Here we describe a unique bidirectional interaction between the host and gut microbiome in sharing precursors for NAD biosynthesis.

Our results also shed light on the metabolism of dietary NAD precursors. The primary dietary precursors for NAD synthesis, amino acids derived from dietary proteins, NAM, and NA are readily absorbed in the upper GI tract, and thus not available to microbes residing in the distal parts of the small intestine and large intestine. Even with a diet free of both NAM and NA, microbial NA production remains high enough to maintain circulating levels in the host. Soluble fiber (traced with ^13^C-inulin) accounts for just under half of NAD synthesis in the large intestine. We find that host-derived NAM almost completely supports microbiome NAD biosynthesis in the small intestine and makes up the other approximately half of NAD synthesis in the large intestine. Microbial NAD generated by any of these routes is turned over to produce NAM and NA that can be taken up by host intestinal tissues and used to regenerate NAD via the Priess-Handler pathway.

The relative importance of NA-dependent (Priess-Handler pathway) NAD synthesis in mammals has been difficult to ascertain. Compared to NAM, NA circulates at a concentration that is nearly an order of magnitude lower and has a much lower turnover flux^21, 46, 59^. The rate limiting enzyme in the Preiss-Handler pathway, *NAPRT,* is expressed in multiple host tissues but the highest enzymatic activity levels are found in the liver and kidney^60^, suggesting that NA might primarily contribute to NAD synthesis in those organs. However, exogenous NA is readily incorporated into NAD in many human cell lines^60, 61^ and some tumors amplify the *NAPRT* gene and depend primarily on the Priess-Handler pathway for NAD^62–64^.

Under conditions of increased NA availability, such as direct supplementation or provision of alternate precursors that can be converted to NA by the microbiome, the Priess-Handler pathway may become more quantitatively important and other NA-dependent mechanisms may also be engaged. An important feature of NAPRT, as compared to NAMPT (NAM-dependent synthesis) is that NAPRT is not subject to feedback inhibition^60^. Incubation of the purified enzymes with NAD results in inhibition of NAMPT^65^, but not NAPRT^66^, and preincubation of cells with NA prevents NAM-dependent NAD synthesis, but not vice versa^67^. Thus, NAD levels can theoretically be increased more by NA than by NAM. In addition, NA at pharmacological doses has lipid-lowering properties and can engage GPR109A, which is responsible for a common side effect - flushing^46, 68^. The mechanism of lipid lowering remains somewhat controversial but may involve inhibition of the diacylglycerol synthesis enzyme DGAT2^69–71^. We find that high dose NR can lead to peak circulating NA concentrations over 50 μM in mice. This concentration is comparable to NA levels in plasma following oral NA intake in humans^42–44^ and thus may be sufficient to transiently activate GPR109A or inhibit DGAT2, mechanisms of action that have not previously been considered relevant for NR. Importantly, if NA-dependent mechanisms play a role in the beneficial effects of NR in mice, it could help explain the lack of translation in some clinical trials - the lower doses used for human studies would be unlikely to support comparable elevation of circulating NA.

We also clarified the metabolic route from NR to NAD. Like Shats et al.^20^, we find that NAD boosting effect of NR depends on microbial deamidation. Our use of NR labeled at multiple different atom types additional distinguishes whether NAD synthesis from NR occurs via NAR or NA. The NAR-dependent route is attractive and would explain how NR could be more effective than either NA or NAM for boosting NAD^48^. Our data, however, conclusively demonstrate that NR contributes to NAD mainly via NA.

NAAD is an effective biomarker for NAD supplementation using NR^8–10, 48^. Although it was originally suggested to result from deamidation of NAD^48^, the discovery that microbes deamidate precursors led to the suggestion that it originates from NA^20^. The labeling patterns in our study strongly support the latter hypothesis, as labeling of NAAD matches that of NA and not NAD.

Our findings suggest some intriguing hypotheses for future investigation. The dependence of microbes on host NAM raises the possibility that the host might modulate microbiome composition or function by limiting luminal NAM transport. In addition, it is unclear whether luminal transport of NAM or microbiome NA production are influenced by stresses such as metabolic syndrome^72–74^. As the colonic microbiome additionally depends on complex carbohydrates that are fermented in the large intestine for NAD synthesis, increased intake of heavily processed food rich in sugar and low in dietary fiber may have a deleterious effect on NAD homeostasis. Such diets are already known to disturb microbiome composition and metabolic activity and are associated with wide range of metabolic diseases in the host^16, 75, 76^. High doses of fructose can also support NAD synthesis in the colon, consistent with recent work showing that fructose can fuel production of microbial metabolites that drive hepatic lipogenesis^25, 77^. It will be important to determine how the cycling of NAD precursors between host and microbiome interacts with host physiology under conditions of metabolic stress.

In conclusion, we have determined the primary precursors used by small and large intestinal microbiota for NAD synthesis, leading to the identification of a vitamin B3 cycle that effectively shares NAD precursors between the host and microbes. Our findings imply that perturbation of NAD metabolism either in the host or microbes has the potential to disrupt NAD homeostasis and impact physiology in the other.

### Experimental models and subject details

All animal procedures were conducted at the University of Pennsylvania and approved by the Institutional Animal Care and Use Committee. 10–12-week-old C57BL/6J male mice were purchased from The Jackson Laboratory and acclimated for at least two weeks with *ad libitum* access to laboratory diet 5010 and water on 12h light: dark cycle (7AM-7PM) before use in experiments. 8-12-week-old germ-free C57BL/6 male mice were obtained from University of Pennsylvania Gnotobiotic Mouse Facility.

### Method details

#### Colonization of germ-free mice

For colonization of germ-free mice (Ex-GF), two adult germ-free mice were co-housed with one special pathogen free (SPF) mouse in each cage for 18 days before tissue collection.

#### In vivo tracing of dietary protein and inulin

^13^C labeled diets were prepared by adding ^13^C-Crude Protein extract from algae (Sigma-Aldrich, 642878) or ^13^C-Inulin (Sigma-Aldrich, 900388) to a premix (modified from normal diet with reduced protein, inulin and starch content, Research Diets Inc., D20030303). The final enrichment for each labeled macronutrient was 25%, and both labeled diets share the same macronutrient composition (63% carbohydrate, 20% protein, 7% fat, 7.5% Inulin, 2.5 % cellulose). 8-week-old male C57BL/6NCrl mice (strain 027; Charles River Laboratories) were first adapted to a composition-matched non-labeled diet for 2 weeks to ensure minimal microbiome perturbation upon diet switch. The non-labeled diet was replaced with a diet that contained a mixture of labeled and unlabeled protein or inulin (1:3) at 9am in the morning, 24 hr before sacrifice, during which mice were given free access to water.

#### In vivo tracing of fructose

Random fed 12–14-week-old C57BL/6J male mice were orally gavaged with 2g/ kg body weight U- ^13^C-Fructose (Cambridge Isotope Laboratories, Tewksbury, MA) dissolved in normal saline. Mice were sacrificed 2h after fructose delivery.

#### In vivo tracing of NA

10 to 12-week-old C57BL/6J male mice were orally gavaged with 1.96 μmoles of 2,4,5,6-^2^H-NA (Cambridge Isotope Laboratories, Tewksbury, MA), a dose equivalent to one-third of dietary intake of NA per day (Manufacturer data show that rodent chow contains 120 ppm nicotinic acid). Blood samples were collected before gavage, 30 min after NA delivery and at the time of sacrifice. Luminal contents and intestinal tissues were collected from the indicated parts of the GI tract.

#### Antibiotic treatment

To deplete gut microbiota mice were given a combination of ampicillin (1 g/L), vancomycin (0.5 g/L), neomycin (1 g/L), metronidazole (0.5 g/L), ciprofloxacin (0.2 g/L), and primoxin (0.5 g/L) in drinking water for 3-5 weeks before use in experiments.

#### NA diet feeding experiments

Custom diets with or without NA were purchased from Research Diets Inc. Cellulose (BW200) and inulin were used as fiber source, and macronutrient and fiber levels were matched to 5010 chow diet. NA diets used to feed germ-free mice were made with 1.5X vitamins and mineral acid casein and subjected to two rounds of gamma-irradiation. 14-week-old C57BL/6J male mice housed in SPF condition were fed the diets with or without NA and given normal drinking water or drinking water with antibiotics cocktail for five weeks. Germ-free and conventional mice were fed with double-irradiated diets for two weeks.

#### *In vivo* tracing of isotopically labeled NAM

2,4,5,6-^2^H-NAM was purchased from Cambridge Isotope Laboratories (Tewksbury, MA). 2,4,5,6-^2^H- NAM tracer was either dissolved in PBS (retro-orbital injection) or normal saline (intravenous infusions). C57BL/6J male mice were retro-orbitally injected with 0.2-5 μmoles of 2,4,5,6-^2^H-NAM in 100μl volume, blood samples were collected from the tail vein either 15min or 2h after injection as indicated in the figure legend and mice were anesthetized to collect blood samples from the portal vein and hepatic vein. Mice were immediately euthanized by cervical dislocation and tissues were collected and clamped in liquid nitrogen.

For intravenous infusion, mice were surgically implanted with a catheter in the right jugular vein and the infusion was performed within a week. NAM tracer infusion was performed as described previously^20^. Briefly, 4mM 2,4,5,6-^2^H-NAM tracer was infused via the jugular catheter at a constant rate of 0.2 nmol/g body weight/min for different period as indicated in the figure legends. Blood samples (∼20μl) were collected from the tail vein using microvette blood collection tubes (Sarstedt, Cat. # 16.440.100). At the end of the infusion period, mice were euthanized by cervical dislocation and tissues were quickly dissected and clamped in liquid nitrogen.

#### FK866 treatment

C57BL6/J or C57BL/6J.Nia male mice given normal drinking water or pre-treated with antibiotics cocktail for four weeks to deplete gut microbiota. Mice were injected with vehicle (45% Propylene glycol, 5% Tween 80, 50% water) or 50 mg/kg FK866 (Selleck Chemicals, Houston, Tx) once (5h NAM infusion) or three times at 8hr intervals (23h NAM infusion). NAM infusion was started 1h after the first FK866 treatment and mice were sacrificed after 5h or 23h.

#### Synthesis of M+9 NR

Scheme 1: Synthesis of silylated M+4 NAM

Scheme 2: Synthesis of silylated M+9 Nicotinamide riboside

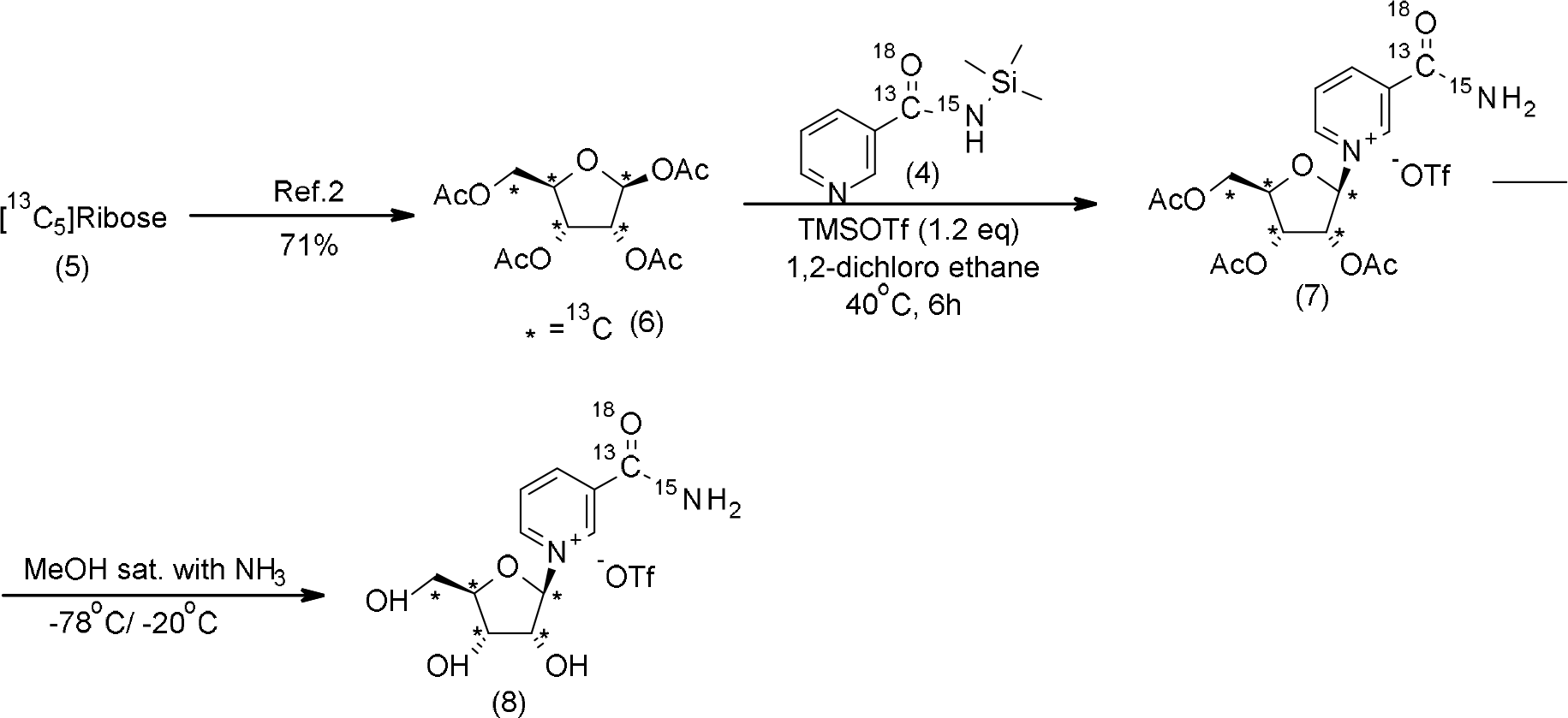

[A] Synthesis of ^13^C, ^18^O and ^15^N labelled M+4 nicotinamide (3, Scheme 1) : Compound (3) was prepared according to reported procedure^20^

[B] Synthesis of Silylated M+4 NAM (4, Scheme 1): A mixture of ^13^C, ^18^O and ^15^N labelled M+4 NAM (0.40 g; 0.032 mol), HMDS (9.00 mL) and TMSCl (0.81 mL, 0.70 g, 0.0064 mol) was heated in an oil bath at 105°C (oil bath temperature) for 20 hours. Upon completion, the reaction mixture was allowed to cool down to room temperature and clear colorless solution was transferred into a 25 mL round-bottom single-neck flask under argon through a cannula and evaporated to dryness on a rotary evaporator followed by drying under high vacuum to give a white crystalline product (0.623 g, 100%).

[C] Synthetic of 1,2,3,5-Tetra-O-acetyl-α/β-D-[^13^C_5_]ribofuranoside ([^13^C_5_]RTA, 6, Scheme 2): Compound (6) was prepared according to reported procedure^78^

[D]General procedure for the synthesis of M+9 NR

*Synthesis of M+9 NR 2,3,5-triacetate (Vorbruggen glycosylation, compound 7, Step-1):* In a flame dried flask under an argon atmosphere, M+4 silylated NAM (0.37g, 0.00187 mol, 1eqv) and M+5 ,^13^C labelled -d-ribose tetraacetate (0.664g, 0.00206 mol, 1.1 eqv) were added to dry 1,2-dichloroethane (12mL). After 5 minutes stirring at rt, TMSOTf (0.40mL, 0.00224 mol, 1.2eqv) was added to the stirred 1,2-dichloroethane solution at the same temperature. Now, the resulting mixture was stirred at 40 °C until complete disappearance of the starting materials. The reaction progress was monitored by ^1^H NMR analysis of the crude mixture. Upon completion, the resulting solution was concentrated, and the yellow oily crude product was used directly for the next step without purification.

*Synthesis of M+9 NR (de-esterification, compound 8, step-4,):* Into a pressure tube closed with a septum, evacuated and filled with argon, [^13^C,^18^O,^15^N]-NRTA -OTf (1.14g, 0.00187 mol, 1eqv) was dissolved in anhydrous methanol (30 mL). The solution was cooled down to -78°C at stirring, and ammonia gas (passed through a tube filled with NaOH) was bubbled into the solution through a long metal needle for ca. 5 min. The reaction solution was additionally stirred at - 78°C for 10 min, and, subsequently, the septum was removed, and the tube was immediately closed with a threaded PTFE cap and transferred into a freezer (−20°C) and kept at -20°C for 6 days. The pressure tube was transferred into an ice bath, and, by using a cannula, the content of the tube was transferred into a recovery flask, cooled down in the same ice bath to 0°C. The recovery flask was attached to a rotary evaporator, and ammonia gas was evaporated without any external heating and immersion in a water bath which resulted in a continuous maintaining of the solution temperature below 0°C. After removal of ammonia, residual methanol was removed at ca. 25°C and an oily residue was kept under high vacuum to give a viscous yellow liquid (1.0g). According to the ^1^H NMR data, the product was contained with the mixture of acetamide, methanol and 2.9 % of M+4 Nam. ^1^H NMR (400 MHz, D2O, δ, ppm): 9.53 (s, 1H, Ar-H), 9.20 (s, 1H, Ar-H), 8.93-8.91 (m, 1H, Ar-H), 8.23-8.19 (m, 1H, Ar-H), 6.18 (d, J = 177Hz, 1H, H-1), 4.66-3.99 (m, 3H, H-2, H-4&H-3), 3.82-3.79 (m, 1H, H-5a), 3.28-3.27 (m, 1H, H-5b); ^13^C NMR (100 MHz; D_2_O,δ, ppm): 170.60 (d, J=17.5Hz, CO), 165.69 (d, J=18.7Hz, CH_3_CONH_2_), 145.62 (d, J= 1.18Hz, C-6_NAM_), 142.60 (C-2_NAM_), 140.34 (C-4_NAM_), 133.40 (d, J= 56.8Hz, C-3_NAM_). 128.33 (d, J= 3.29Hz, C-5_NAM_), 99.91 (dd, J= 40.02 Hz &3.08Hz, C-1_ribose_), 87.67 (t, J= 39.9 Hz, C-4_ribose_), 77.42 (t, J=38.32 Hz, C-2_ribose_), 69.75 (t, J= 36.19 Hz, C-3_ribose_), 60.17 (d, J= 41.06 Hz, C-5_ribose_); ^19^F NMR (377 MHz, D_2_O) δF: −78.82 (s); HRMS calcd for ^13^C_6_ ^12^C_5_H_15_ ^15^N^14^N^18^O^16^O_4_ [M]^+^ 264.1189found 264.1194.

#### NR oral gavage

For NR experiments mice were orally gavaged with NR (500mg/kg body weight) or a mixture of custom-synthesized M+9 NR and unlabeled NR at a ratio of 1:2.6 in PBS (600 mg/kg body weight). Blood samples were collected at different time points. Three hours after oral dosing, mice were anaesthetized to collect blood samples from hepatic and tail veins and sacrificed by cervical dislocation to collect tissues. In cases where the tracer was diluted with unlabeled molecules, the ratio was verified by mass spectrometry and used to correct for the total contribution of the administered molecule to a given metabolite of interest. This method is expected to be accurate for following any single molecule, but could underestimate the frequency of two labeled molecules coming together by chance, such as regeneration of labeled NR or NAR in the intestinal lumen from labeled ribose and NAM or NA.

#### Metabolite extraction from serum and tissue samples

Serum samples were thawed on wet ice before metabolite extraction. 50ul of 100% methanol was added to 5ul serum, vortexed and incubated on dry ice for 10 minutes, and centrifuged at 16,000 g for 20 minutes, and 10 ul of water was added to 40ul of serum extracts and use for LC-MS analysis. Frozen tissues were weighed and ground with liquid nitrogen in a mortar and pestle or cryomill (Retsch). Cryomilled tissues were extracted with 40:40:20 acetonitrile:methanol:water (25 mg tissue / ml extraction buffer), vortexed and incubated on ice for 10 min. Tissue samples were then centrifuged at 16,000 g for 30 minutes. The tissue supernatants were transferred to new Eppendorf tubes and then centrifuged again at 16,000 g for 25 minutes to remove and residual debris before LC-MS analysis.

#### Metabolite Measurements

Extracts were analyzed within 36 hours by liquid chromatography coupled to a mass spectrometer (LC-MS). The LC–MS method involved hydrophilic interaction chromatography (HILIC) coupled to the Q Exactive PLUS mass spectrometer (Thermo Scientific) as reported previously^79^. The LC separation was performed on a XBridge BEH Amide column (150 mm 3 2.1 mm, 2.5 mm particle size, Waters, Milford, MA). Solvent A is 95%: 5% H2O: acetonitrile with 20 mM ammonium bicarbonate, and solvent B is acetonitrile. The gradient was 0 min, 90% B; 2 min, 90% B; 3 min, 75%; 7 min, 75% B; 8 min, 70%, 9 min, 70% B; 10 min, 50% B; 12 min, 50% B; 13 min, 25% B; 14 min, 25% B; 16 min, 0% B, 20.5 min, 0% B; 21 min, 90% B; 25 min, 90% B. Other LC parameters are: flow rate 150 μl/min, column temperature 25°C, injection volume 10 μL and autosampler temperature was 5°C. The mass spectrometer was operated in both negative and positive ion mode for the detection of metabolites. Other MS parameters are: resolution of 140,000 at m/z 200, automatic gain control (AGC) target at 3e6, maximum injection time of 200 ms and scan range of m/z 75-1000. Raw LC/MS data were converted to mzXML format using the command line “msconvert” utility^80^. Data were analyzed via the MAVEN software using in-house metabolite library established from authentic standards.

*Nicotinate* was detected with a 13-min method using the same LC buffers and column but a different gradient at a flow rate of 300 μl/min: 0 min, 90% B; 2 min, 90% B; 5 min, 0 %B; 8 min, 0 % B; 9 min, 90% B; 13 min, 90% B. A SIM scan was used covering m/z 120 – 130 in positive mode.

*MS2 of unlabeled and labeled NAD*: targeted MS2 was performed using the PRM function with the parent ions at m/z 664.1, 669.1, 674.1 and NCE 20%. Other parameters are, resolution 17500, AGC target 5e5, maximum injection time 200 ms, isolation window 1.5 m/z. LC condition was the same as the above 25-min method for water soluble metabolites.

#### Statistical analysis

Unless otherwise noted, data are presented as mean ± SEM. One-way or two-way ANOVA was used with Tukey’s post-hoc test for comparisons of 3 or more groups. Student’s t-test was used for 2 group comparisons or to determine nominal significance. Asterisks displayed in the figure denote statistical significance (p < 0.05; **, p < 0.01; ***, p<0.001).

## Acknowledgments

This work was funded and supported by a Crohn’s and Colitis Career Development Award to KC; the Howard Hughes Medical Institute and Burroughs Wellcome Fund via the PDEP and Hanna H. Gray Fellow Programs awarded to MRM; NIH grants CA211437 to WL, DP1DK113643 to JDR, R01DK098656 and R01AG043483 to JAB. We acknowledge support from the Regional Metabolomics and Fluxomics Core and the Rodent Metabolic Phenotyping Core of the Penn Diabetes Research Center P30-DK19525 and S10-OD025098, as well as the CINJ Cancer Center Support Grant, Rutgers Cancer Institute of New Jersey Metabolomics Shared Resource, supported, in part, with funding from NCI-CCSG P30CA072720-5923.

## Author Contributions

KC, MRM, JDR, and JAB conceived and designed the project. KC and MRM performed and analyzed most experiments. WL, XZ, YRB, SRL, HCD, TC, LJ, and CJ performed specific *in vivo* experiments and analyses. MM, FH, and MEM designed and synthesized M+9 NR. KC and JAB wrote the manuscript with input from all authors.

---

**Supplementary Figure 1.**
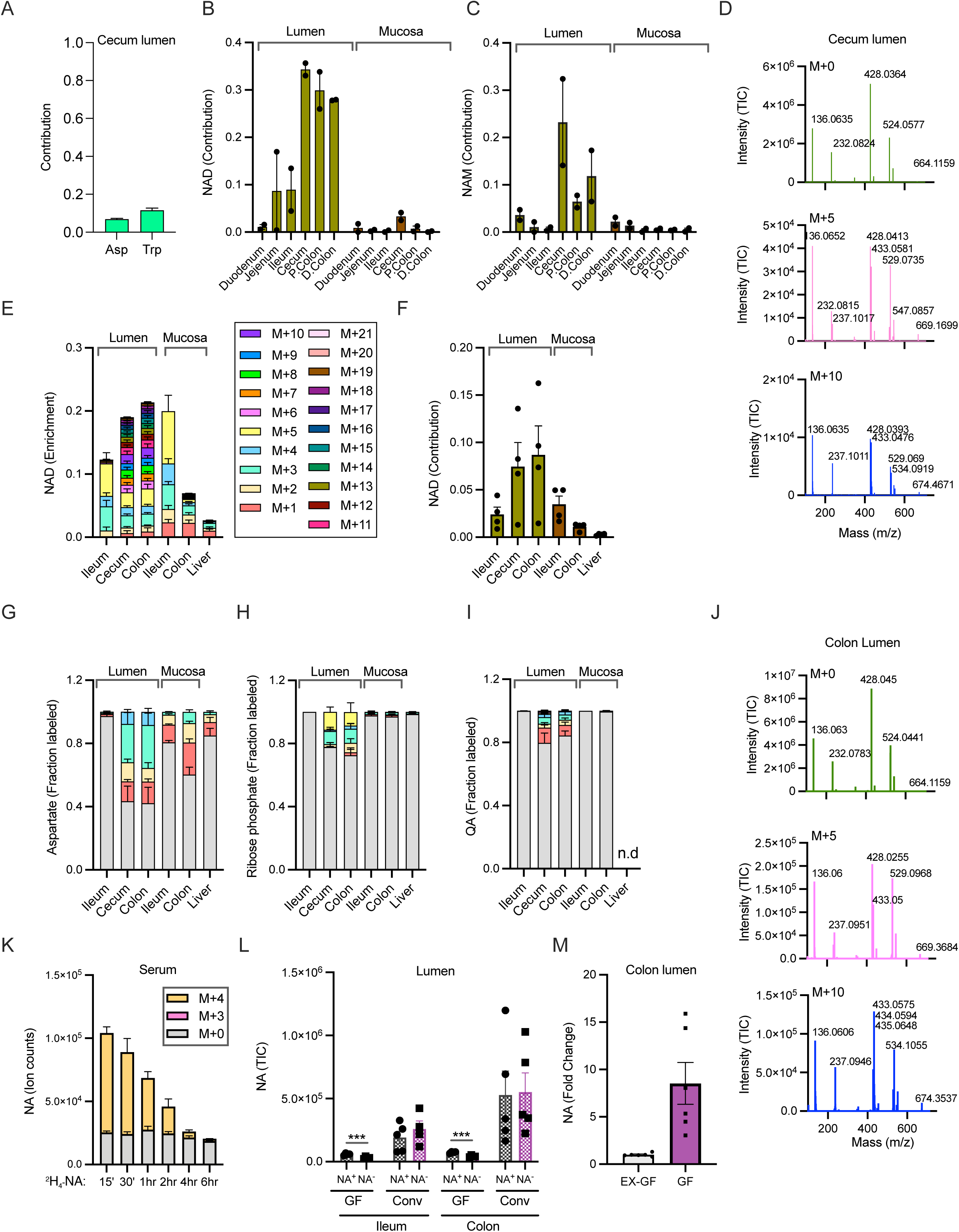
Site-specific use of dietary NAD precursors by microbes along the GI tract. A. Contribution of U-^13^C-protein to aspartate and tryptophan levels in the cecum lumen of mice fed with labeled protein diet. n=3 mice. B. Contribution of U-^13^C-inulin to NAD synthesis in different parts of gastrointestinal tract of mice fed with labeled protein. n=2 mice. C. Contribution of U-^13^C-inulin to nicotinamide in the gut lumen of mice treated as in B. n=2 mice. D. MS^2^ fragments of NAD detected in the cecum lumen of mice fed with U-^13^C-inulin fed as in B. E-F. Enrichment of NAD labeling (E) and contribution to NAD synthesis (F) in mice orally gavaged with U-^13^C-fructose for 2h at a dose of 2g/kg. n=4 mice. G-I. Enrichment of aspartate (G) ribose phosphate (H) and quinolinic acid (I) labeling in mice treated as in E. J. MS^2^ fragments of NAD detected in the colonic lumen of mice treated as in E. K. Labeled and unlabeled NA in the serum after oral gavage of 1.96 μmoles of [2,4,5,6-^2^H]-NA n=2- 6. L. NA content in the lumen of germ free (GF) and conventional (Conv) mice fed diet with (NA^+^) or without nicotinic acid (NA^-^). n=4-5 mice per group. M. Abundance of NA in the colonic lumen of germ-free (GF) and Ex-germ free (Ex-GF) mice colonized with microbiota from SPF mice. n=4 per group.

**Supplementary Figure 2.**
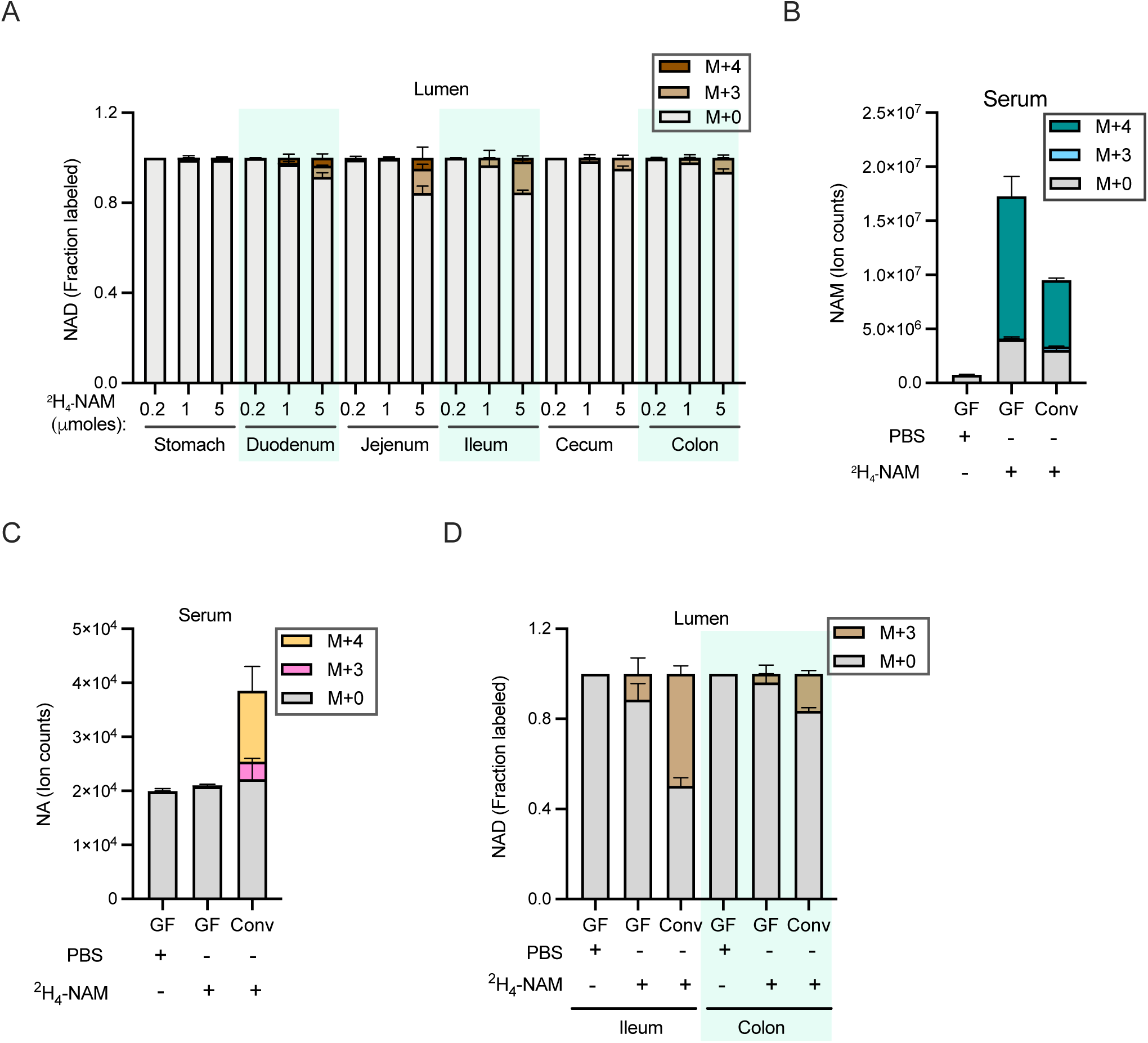
Circulating NAM labels luminal NAD. A. Fraction labeled NAD in luminal contents collected from mice retro-orbitally injected with 5 μmoles [2,4,5,6-^2^H]-NAM and sacrificed after 15 min. n=2-3 mice per group. B. NAM levels in the serum samples collected from germ-free (GF) and conventional (Conv) mice 2h after retro-orbitally injection of either PBS or 5 μmoles of [2,4,5,6-^2^H]-NAM. n=4 mice per group. C. Serum NA labelling in mice treated as in B. D. Fraction labeled NAD in the luminal samples collected from mice treated as in B.

**Supplementary Figure 3.**
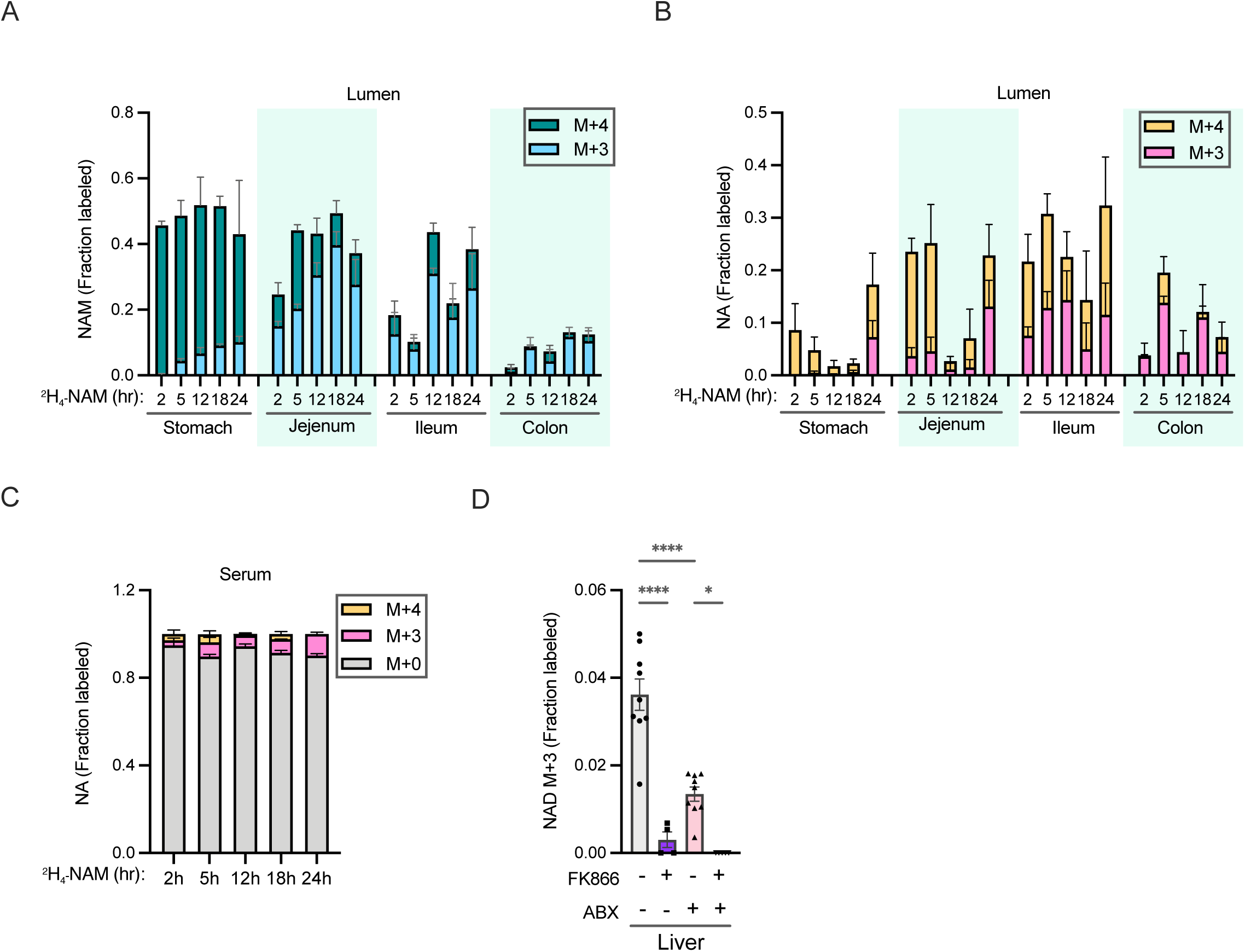
The gut microbiome provides metabolic flexibility to bypass salvage synthesis in liver. A. Fraction isotope labeling of NAM in the lumen of mice intravenously infused with 4mM [2,4,5,6-^2^H]-NAM. n=2-4 mice per group. B. Fraction isotope labeling of NA generated by microbial deamidation of [2,4,5,6-^2^H]-NAM in mice treated as in A. C. Labeling of serum NA in mice infused as in A. D. Fraction labeled NAD in the liver from control and antibiotics (ABX) treated mice intraperitoneally injected with vehicle or FK866 and infused with [2,4,5,6-^2^H]-NAM for 5h. n=3-5 mice per treatment group.

**Supplementary Figure 4.**
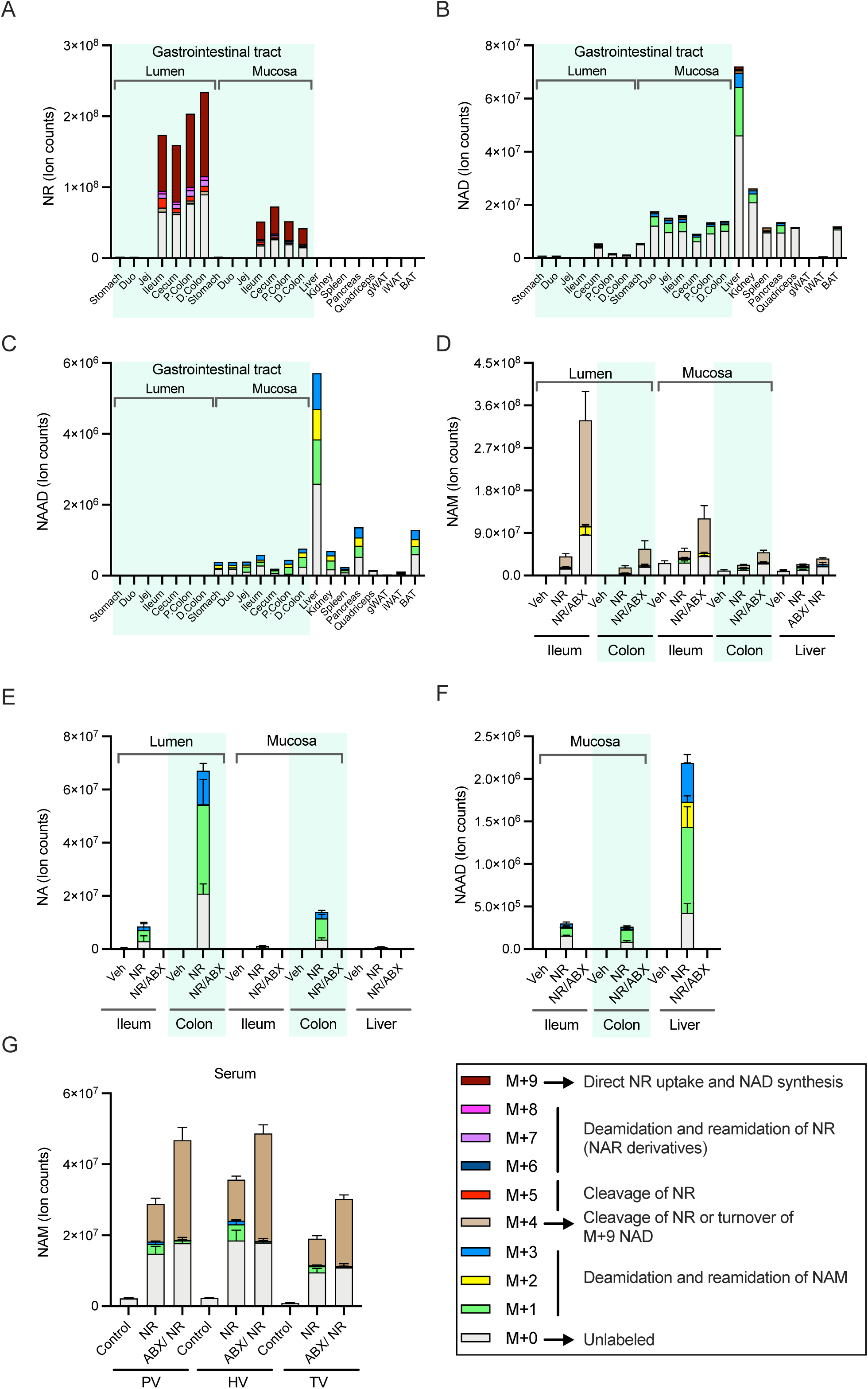
The gut microbiome is required to generate deamidated precursors from orally delivered NR. A-C. NR (A), NAD (B), and NAAD (C) labeling in the lumen and tissues of a mouse orally gavaged with NR at a dose of 600 mg/kg body weight) for 3h. D-F. NAM (D), NA (E), and NAAD (F) labeling in the lumen and tissues of mice orally gavaged with mixture of unlabeled NR at a dose of 600 mg/kg body weight for 3h. n=4-5 mice per treatment group. H. Serum NAM levels in mice treated as in D.

